# The Delta variant SARS-CoV-2 spike protein uniquely promotes aggregation of pseudotyped viral particles

**DOI:** 10.1101/2022.04.07.487415

**Authors:** Jennifer D. Petersen, Jianming Lu, Wendy Fitzgerald, Fei Zhou, Paul S. Blank, Doreen Matthies, Joshua Zimmerberg

## Abstract

Individuals infected with the SARS-CoV-2 Delta variant, lineage B.1.617.2, exhibit faster initial infection with a higher viral load than prior variants, and pseudotyped particles bearing the SARS-CoV-2 Delta variant spike protein induce a faster initial infection rate of target cells compared to those bearing other SARS-CoV-2 variant spikes. Here, we show that pseudotyped particles bearing the Delta variant spike form unique aggregates, as evidenced by negative stain and cryogenic electron microscopy (EM), flow cytometry, and nanoparticle tracking analysis. Viral particles pseudotyped with other SARS-CoV-2 spike variants do not show aggregation by any of these criteria. The contribution to infection kinetics of the Delta spike’s unique property to aggregate is discussed with respect to recent evidence for collective infection by other viruses. Irrespective of this intriguing possibility, spike-dependent aggregation is a new functional parameter of spike-expressing viral particles to evaluate in future spike protein variants.

## Introduction

Genetic variants of the severe acute respiratory syndrome coronavirus 2, SARS-CoV-2, continue to evolve as the virus circulates worldwide, and each variant holds the potential to evade acquired immunity and re-ignite the COVID-19 pandemic [1]. SARS-CoV-2 is an enveloped RNA virus containing a single stranded, positive-sense genome [2]. Decorating the surface of the virus is the prominent, club-shaped spike glycoprotein (spike) which drives fusion between the viral envelope and host cell membranes to deliver the viral genome [3,4]. The spike is highly immunogenic, eliciting a robust neutralizing antibody response, and is the immunogen encoded by the highly efficacious mRNA vaccines [5,6]. Due to its essential role in viral entry and immunity, mutations occurring on the spike require careful genetic, structural, and functional surveillance.

The fully assembled, prefusion spike consists of a trimer of spike protomers, each of which is highly glycosylated [7,8]. Proteolytic cleavage of the SARS-CoV-2 spike by furin during biosynthesis nicks the spike into two subunits, S1 and S2 [5]. The S1 subunit contains the receptor binding domain (RBD) which can be in an up (receptor accessible) or down (receptor inaccessible) conformation [6], the N-terminal domain (NTD), and two C-termini. S1 caps the S2 subunit which harbors the membrane fusion machinery [5,6,9]. The spike is highly flexible due to a hinged stalk [10,11], which may facilitate RBD binding to the host cell receptor, angiotensin converting enzyme II (ACE2) [5,9,12]. Once attached to the host cell by ACE2 binding, a second cleavage event occurs in S2 by a host cell protease, either TMPRSS2 at the cell surface or cathepsin in the endosomal membrane, depending on cell type, protease availability, and viral variant [13,14]. Upon S2 cleavage, the fusion peptide undergoes large conformational changes that drive fusion between membranes [15,16].

Millions of SARS-CoV-2 genomic sequences have been cataloged since the original strain emerged in Wuhan, China in December 2019 [17,18]. Amino acid substitutions or deletions in the spike that impart fitness benefits to the virus have evolved and sometimes converged independently in different geographical locations [19,20] giving rise to several variants designated Variants of Concern (VOCs) by the World Health Organization [20,21]. The first was a single amino acid change from an aspartic acid to a glycine at amino acid 614 (D614G), that increased transmissibility [22] possibly by stabilizing the prefusion spike trimer [23], increasing the RBD up/receptor accessible conformation rate [24], and increasing the spike density on the virion [25]. This D614G variant, designated Pango lineage B.1 [26], rapidly supplanted the original Wuhan strain globally [27] by May 2020, and the D614G substitution is present in all subsequent variants [28]. Next, the VOC Alpha (lineage B.1.1.7) emerged in the UK and became dominant worldwide by early 2021, owing to additional substitution mutations occurring in the spike RBD that increased ACE2 receptor affinity, deletions in the NTD that increased immune escape, and a mutation in S2 that may enhance membrane fusion potential [29–31], collectively leading to about two-fold enhanced transmissibility of Alpha compared to other contemporaneously circulating variants [32].

The Alpha variant prevailed until it was outcompeted by the VOC Delta (lineage B.1.617.2), a member of the B.1.617 lineage, that arose in India and became globally dominant in mid-2021 [33]. The Delta variant was considerably more transmissible than Alpha [34], infected individuals faster, produced earlier detection by PCR test with higher viral load [35–37], and was more pathogenic [38]. The Delta variant’s spike displayed a different combination of mutations compared to Alpha, that were predicted to reduce sensitivity to neutralizing antibodies [39–41] and increase ACE2 receptor binding, spike stability, viral infectivity, fusogenicity, and pathogenicity [42–44].

Functional consequences of spike mutations can be tested using engineered mimics of enveloped viruses, called pseudotyped particles (PPs). PPs are produced by co-transfecting producer cells with plasmids encoding a capsid protein from a parental virus (typically a retrovirus or arbovirus capsid), the spike glycoprotein of interest, and a reporter gene that produces a fluorescent or luminescent protein signal upon host cell entry. The capsid core buds efficiently from the producer cell while incorporating the heterologous spike and encapsulating the reporter gene [45,46].

Recently, studies comparing the entry rate of PPs packaged with various SARS-CoV-2 spike variants into target cells showed that PPs bearing the Delta spike (Delta PPs) drove markedly faster initial infection, and greater infection overall, than other variants [35,43]. This phenotype is similar to real-world reports of faster initial infection by Delta SARS-CoV-2 [37].

To search for an ultrastructural correlate to the increased initial infection rate of Delta PPs, the structures of murine leukemia virus (MLV)-based PPs bearing several different SARS-CoV-2 spike variants were examined by negative stain transmission electron microscopy (TEM): D614G, Alpha, Delta, Delta sublineage AY.4.2 (Delta AY.4.2), and Omicron sublineage BA.1 (Omicron BA.1) (lineage B.1.1.529), along with ‘Bald PPs’, that lacked a spike glycoprotein. Negative stain TEM showed that PPs bearing Delta and Delta AY.4.2 spikes uniquely clustered into aggregates, while the other variant spike PPs and Bald PPs did not. There are clear indications of spike-spike tip interactions mediating this clustering. The aggregation of Delta and Delta AY.4.2 PPs was confirmed by flow cytometry and nanoparticle tracking analysis (NTA). Cryo-electron microscopy of Delta PP aggregates showed a variety of distances between the envelopes of PP, many consistent with the spike-to-spike interactions seen in negative staining. Implications for Delta spike-mediated aggregation for the kinetics of initial viral infection are discussed.

## 2. Materials and Methods

### 2.1 Cell culture and Production of SARS-CoV-2 Variant Spike PPs

HEK 293T cells were a gift from the laboratory of Dr. Gary Whittaker (Cornell University). Cells were maintained in DMEM (Genesee Scientific Corporation, Cat #: 25-500) with 10 % FBS (Corning, Cat#35-010-CV), 1X Pen-Strep (Thermo Fisher, Cat#15140-122) and 250 mg/ml G418 (TOKU-E, Cat SKUG001). The day before transfection, 8 million HEK 293T cells were plated on a 10-cm cell culture dish (EZ BioResearch, Cat# C3100) in 16 ml of DMEM with 10% FBS (no antibiotics were added). Three plasmids pCMV-MLV-gag-pol, pTG-Luc and SARS2-COV2 spike expression vector were transfected into HEK 293T cells at the ratio of 3:4:3 using Lipofectamine 3000 transfection reagent (ThermoFisher Scientific, Waltham, MA). Accession IDs of the spike proteins used in the study were: D614G (B.1), EPI_ISL_464996; Alpha (B.1.1.7) EPI_ISL_736724; Delta (B.1.617.2) EPI_ISL_2020954; Delta AY.4.2, EPI_ISL_5423152; Omicron BA.1 (B.1.1.529) EPI_ISL_6699757. To increase expression of the spike protein on the cell surface, the C-terminal 19 amino acids were deleted [47,48]. 48 hours post-transfection, the supernatants were collected and centrifuged at 290 g for 7 min at 4°C. The supernatants were then passed through a 0.45 μm syringe-tip filter [45] and stored on ice or 4°C until use.

### 2.2 Negative stain transmission electron microscopy (TEM) and immunogold labeling

PPs were stored on ice after harvest from producer cells and processed as follows for TEM within 4 hours of harvest or after overnight storage at 4°C. All reagents were obtained from Electron Microscopy Sciences (EMS, Hatfield, PA) and all steps took place at room temperature unless otherwise specified. PPs, suspended by gentle trituration, were adhered to freshly glow discharged, formvar and carbon-coated, 300-mesh gold EM grids (EMS) by inverting grids for 2 min on 5 μl drops of culture medium supernatant containing PPs. Grids were rinsed by transferring the grids across 3 droplets of filtered DPBS (Dulbecco’s phosphate buffered saline, Mg and Ca-free, Gibco, Paisley, UK). All solutions were filtered through a 0.22 μm syringe filter with RC membrane (EMS) before contact with the EM grid. For negative staining, grids were rinsed twice with filtered, distilled water and then placed on a drop of filtered 1% aqueous uranyl acetate for 1 minute, and then blotted to dryness with filter paper (Whatman #1).

To immunogold label the spike protein S1 subunit, grids with adhered PPs were floated on drops of blocking solution containing 2% BSA (Sigma, St. Louis, MO) in DPBS for 10 min. Primary antibody to SARS-CoV-2 spike subunit 1 (Sino Biological, Beijing, Cat. No. 40589-T62), was diluted 1:25 in blocking solution and grids were transferred to drops of primary antibody for 30 minutes. Then, grids were transferred across two drops of blocking solution for 10 minutes before incubation with 10 nm gold-conjugated donkey-α-rabbit secondary antibody diluted 1:20 in blocking solution for 30 min. Grids were covered during incubation steps to prevent evaporation. Finally, grids were rinsed with 3 drops of DPBS and negative stained as described above. Grids were observed using a FEI Tecnai T20 transmission electron microscope (Thermo Fisher, Waltham, MA) operated at 200 kV, and images were acquired using a NanoSprint1200 CMOS detector (AMT Imaging, Woburn MA). Images were prepared for display using Photoshop 2022 (Adobe). The sizes of the Delta variant PP aggregates were measured using Fiji. A freehand perimeter was traced around each aggregate to obtain area and size measurements.

### 2.3 Flow cytometry of PPs

PPs at 4 hours, 24 hours and 7 days were labelled with 0.5 μM BODIPY FL (Thermo Fisher, Waltham, MA), a hydrophobic dye that stains lipids, in PBS for 15 minutes at 4°C. Samples were acquired on a BD Symphony A5 flow cytometer (BD Biosciences, Franklin Lakes, NJ) with forward and side scatter in log mode and thresholding set on BODIPY-positive events. MegaMix-Plus SSC microparticles (Diagnostica Stago, Parsippany, NJ) were used to optimize scatter settings and establish size regions. Data was analysed in FlowJo 10.8 (BD Biosciences, San Jose, CA).

### 2.4 Nanoparticle tracking analysis (NTA) of PPs

The size and concentration of PPs at 4 hours, 24 hours, and 7 days were characterized by NTA using a NanoSight NS300 (Malvern Instruments Ltd, Malvern, UK) equipped with a 405-nm laser. PPs were diluted 1:100 in PBS, loaded into 1 ml syringes, introduced into the sample chamber using a syringe pump and three 60 second videos were recorded per sample. Acquisition and analysis settings were kept constant for all samples. Size distributions and concentrations were analyzed in NTA software, version 3.4. Aggregate percentages were calculated by summing the area under the size distribution curve for all species greater than the monomer, normalized by the total area under the curve.

### 2.5 Cryo Electron Microscopy

7 μl of PPs at 4 or 24 hours were adhered to the top side of freshly glow discharged, 200-mesh Quantifoil gold R2/1 EM grids (EMS), back-blotted for 6-8 s after a 30-s pre-blotting time and immediately plunge frozen into liquid ethane using a GP-EM2 (Leica Microsystems, Wetzlar, Germany) with a chamber maintained at 20 °C and 95% humidity. Grids were screened using a FEI Tecnai T20 transmission electron microscope (Thermo Fisher, Waltham, MA) operated at 200 kV, and images were acquired using SerialEM [49] and direct electron detector K2 Summit (Gatan Inc., Pleasanton, CA) at a nominal magnification of 25,000x at a binned pixel size of 3.04 Å/px and about −3 μm defocus. At a dose rate of 8 e^-^/px/sec, 10 s exposures with 0.2 s frames were collected. Each 50-frame-movie was motion corrected using SerialEM. To detect the size of the larger Delta variant PP aggregates, a medium magnification of 1700x and a binned pixel size of 88.7 Å/px was used.

## 3. Results

### 3.1 PPs bearing the Delta spike variant cluster into aggregates

Negative stain TEM was conducted to compare the structure of Delta PPs to those bearing other SARS-CoV-2 spike variants: D614G, Alpha, Delta AY.4.2, and Omicron BA.1. As a control, PPs produced in the absence of a spike, referred to as “Bald”, were also examined. For each set of experiments, PPs were produced in parallel, maintained on ice or at 4°C, handled under identical conditions, and prepared for TEM within 4 hours of harvest. By negative stain TEM, all spike variant PPs showed a population of ~120 nm diameter spherical particles (Figure 1 A-D), the expected size of MLV-based particles [50]. PPs were abundant in the culture medium without concentration, producing a density on the surface of the EM grid of about five PPs per 60 μm^2^ of EM grid surface. Also uniformly distributed on the grid surface were ~25 nm lipoprotein-like particles, sometimes closely associated with PPs, including the Bald PPs. Collapsed dehydrated vesicles resembling extracellular vesicles were observed occasionally, but rarely.

**Figure 1.**
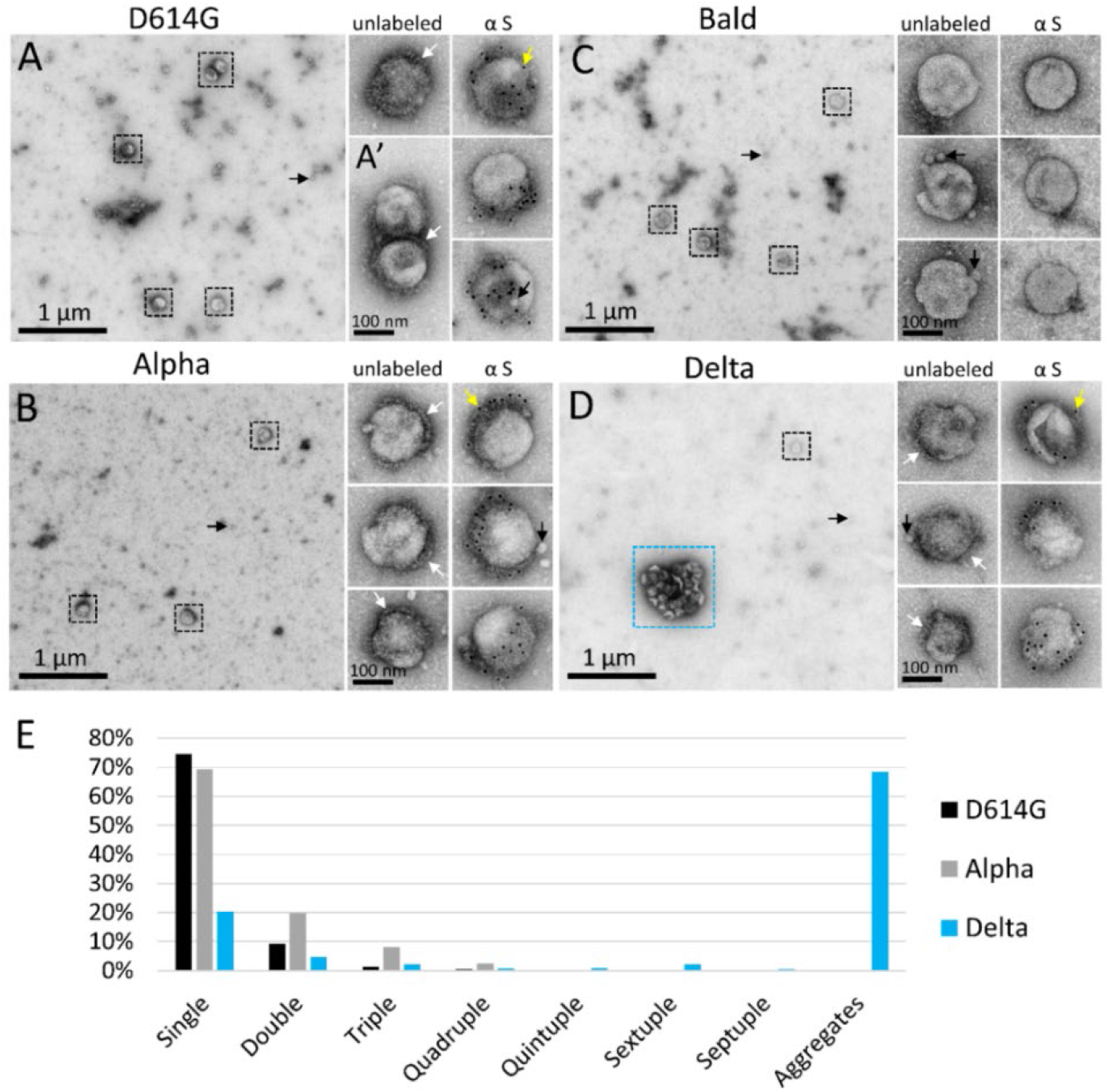
Negative stain TEM detects unique aggregation of PPs bearing the Delta spike. (A-D) Low magnification negative stain TEM overviews of PPs bearing SARS-CoV-2 spike variants (A) D614G, (B) Alpha, (C) Bald, or (D) Delta. Black boxes surround individual PPs; blue box indicates an aggregate of Delta PPs. To the right are enlarged views of individual unlabeled PPs and PPs immunogold labeled for spike protein. (A’) Example of a doublet of D614G PPs. White arrowheads show areas on PPs where a fringe of spike proteins is clearly visible. Yellow arrowheads indicate 10 nm immunogold particles labeling the spike S1 subunit, which appear as black dots. Small black arrows indicate lipoprotein-like particles. (E) Frequency of aggregates of eight or more PPs per variant.

Spike variant PPs often displayed a fringe of spike-like projections on a portion of their surface. Immunogold labeling, directed against the S1 subunit, labeled spikes on D614G, Alpha, Delta, and Delta AY.4.2 variant PPs, but did not label the Omicron BA.1 PPs. The negative control Bald PPs that lacked spikes appeared smooth compared to the others and did not immunogold label (Figure 1).

While the different spike variant PPs looked similar on an individual basis, there was a dramatic difference in the interaction between Delta and Delta AY.4.2 PPs, versus the other variant and Bald PPs. The vast majority of D614G, Alpha, Omicron BA.1, and Bald PPs existed as single PPs or in groups of 2-3 closely associated PPs (Figure 1A’). In contrast, in three experimental replicates, half or more of the Delta PPs were sequestered into aggregates of eight or more (up to dozens) PPs (Figure 1D and Figure 2). To quantify the degree of aggregation of the variant PPs, low magnification images were collected at random, and PPs counted and categorized as single PPs spaced more than 60 nm apart (beyond the length of two spike proteins), or in a cluster of two, three or more PPs. In three experimental replicates, D614G and Alpha PPs were present as single PPs about 75% of the time. Omicron BA.1 PPs (two experimental replicates) were also primarily present as single PPs (not quantified). Bald PPs aggregated at the same rate as D614G and Alpha, suggesting that these spike variants had a comparable and minimal effect on PP interactions that promote aggregation.

**Figure 2.**
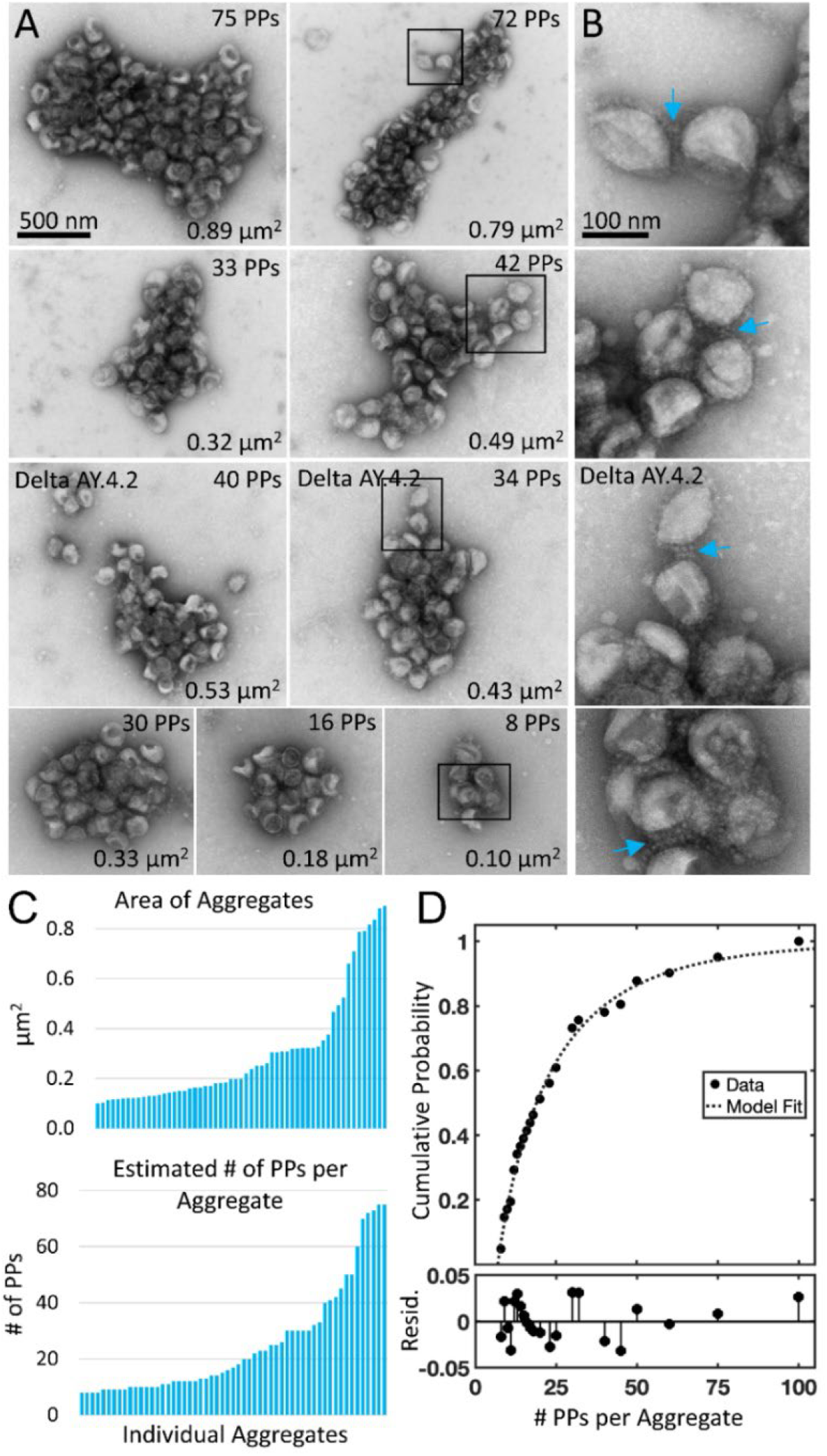
Negative stain TEM of Delta and Delta AY.4.2 aggregates observed 4 hours after harvest from producer cells. (A) Delta and Delta AY.4.2 PP aggregates representing range of sizes at the 4-hour timepoint. The number of PPs estimated per aggregate is indicated in the upper right corner of each image and the area occupied by the aggregate indicated in the lower right corner. Two Delta AY.4.2 PP aggregates are labeled, all others shown are Delta PPs (unlabeled). (B) Areas boxed in A are shown enlarged to the right of each image. Blue arrows highlight spike tip interactions occurring between PPs at the periphery of aggregates. (C) Graphs show ordered lists of the estimated number of PPs and areas of aggregates. (D) Truncated Weibull Distribution for Delta Variant. Negative stain data at 4 hours was fit by the general model: f(x) = (wblcdf(x,a,b)-wblcdf(7,a,b))/(1-wblcdf(7,a,b)) where wblcdf is the Weibull Distribution and x is the PPs per aggregate, 7 is the truncation value, and a and b are the Weibull scale and shape parameters. Coefficients (with 95% confidence bounds) are a = 11.47 (8.35, 14.60) and b = 0.68 (0.57, 0.79). Goodness of fit measure Sum of Squared Error (SSE) = 0.009 and Standard Error of Regression (RMSE) = 0.021, and Resid. are the residuals or differences between the model fit and the data.

The average Delta PP aggregate occupied 0.3 μm^2^ and was 700 nm in its longest axis (57 Delta PP aggregates containing eight or more PPs measured in three experimental replicates, average area 0.30 ± 0.23 μm^2^, average major axis 708 ± 300 nm) (Figure 1D and Figure 2). The number of PPs in an average sized aggregate was ~25 PPs, however this is likely an underestimate as PPs inside the aggregate were obscured from view, particularly in the case of larger aggregates. The Delta variant sublineage Delta AY.4.2 exhibited the same aggregation property (one experimental replicate, 48 % of Delta AY.4.2 PPs in aggregates of 8 PPs or more).

To gain insights into the interactions between Delta PPs in aggregates, PPs on the periphery of aggregates were examined at high magnification. Examples of Delta PPs with clear indications of spike tip interactions were observed between PPs associated with the edge of aggregates (Figure 2C, white arrows). However, since negative stain requires the PPs to be dried in a thin layer of electron dense stain, it was not possible to see into deeper areas of the aggregate where PPs were layered on top of each other.

From this dataset, the aggregation properties of Delta are different from Alpha and D614G. This conclusion is supported by several lines of evidence. First, whereas Delta exhibits overdispersion, Alpha and D614G exhibit underdispersion. Second, whereas the Alpha and D614G have similar distributions with low probability of aggregates with 3 or more PPs, Delta has a long tail with higher probability of aggregates with 3 or more PPs (> 10x). Third, Alpha and D614G distributions are described by truncated Poisson distributions with similar parameter while the Delta distribution is better described by a truncated Weibull distribution (Figure 2D).

### 3.2 Delta spike variant PPs continue to aggregate in solution

The same samples of PPs that were prepared within 4 hours of harvest, were stored at 4°C overnight (still suspended in culture medium) and prepared again for negative stain TEM the next day. PPs of D614G, Alpha, Omicron BA.1, and Bald PPs remained predominantly singles and showed minimal aggregation. However, after overnight storage, the aggregates of Delta and Delta AY.4.2 PPs were much larger, less frequent, and required lengthy searching with the microscope to find. In parallel, there was a concomitant reduction in single Delta and Delta AY.4.2 PPs on the grid surface indicating that single PPs continued to become sequestered in aggregates while diffusing in the culture medium and aggregates themselves likely aggregated. Aggregates at 24 hours were irregularly shaped, sometimes donut-shaped, and up to several microns in size (Figure 3).

**Figure 3.**
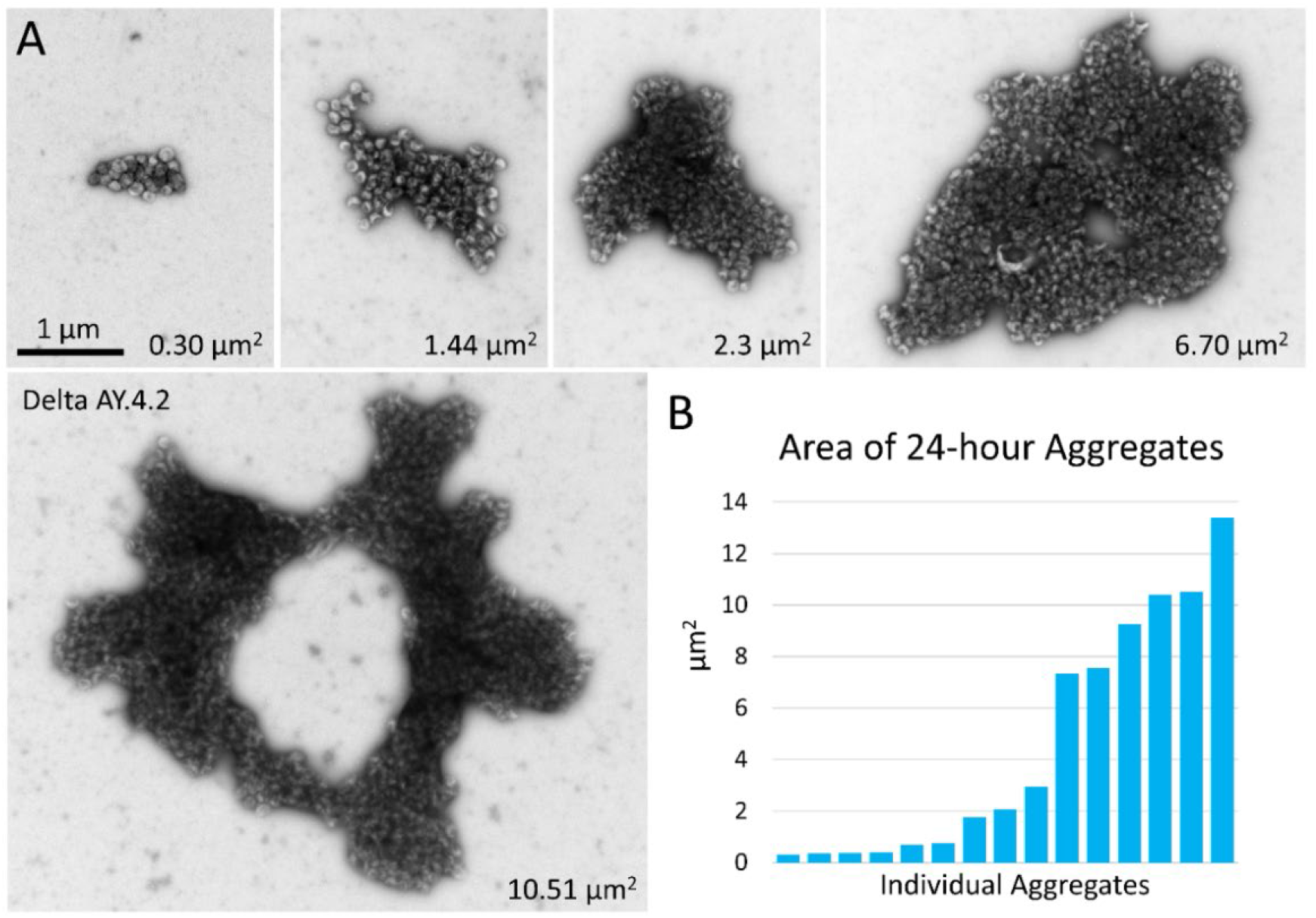
Negative stain TEM of Delta and Delta AY.4.2 aggregates 24 hours after harvest from producer cells. (A) Images of aggregates of Delta and Delta AY.4.2 PPs representing range of sizes observed after storage overnight at 4°C. The area occupied by each aggregate is indicated in the lower right corner. One donut-shaped Delta AY.4.2 aggregate is labeled, all other aggregates are Delta PPs (unlabeled). All images are scaled the same and scale bar is indicated. (B) Graph shows an ordered list of the areas of Delta aggregates.

### 3.3 Flow cytometry and NTA confirm population of aggregated Delta Spike variant PPs

Since negative stain TEM permits observation of only those entities that adhere to the surface of the EM grid, we applied two alternative methods, flow cytometry and nanoparticle tracking analysis (NTA) which sample particles present in the entire solution volume, to corroborate the negative stain TEM observations. All variant PPs measured by flow cytometry and NTA within four hours of harvest from producer cells detected particles predominantly in the single PP range (not shown, n=2). However, after 24 hours stored at 4°C, flow cytometry and NTA of PPs detected a significant population of larger particles in the Delta PPs that was not present in the other variant PPs, or the Bald PPs (Figure 4A and C, n=2). Because NTA does not count aggregates above 1 μm in size, the larger aggregates observed by negative stain TEM are not reflected in the NTA analysis. However, the abundance of single Delta PPs (peak ~150 nm), reduced by half compared to other PPs, is likely explained by their sequestration into aggregates (Figure 4C). Despite these differences in sensitivity, a significant population of Delta PPs were observed in large aggregates by flow cytometry and NTA, that were absent in PPs bearing other variant spikes and Bald PPs. After storage of PPs at 4°C for 7 days, flow cytometry was repeated and still minimal aggregation of D614G, Alpha, or Bald PPs was detected, suggesting that these variants and Bald PPs have less tendency to aggregate under these conditions (Figure 4B, n=2). However, after 7 days there was evidence for some deterioration in the functionality of PP samples, as cell entry was very low (data not shown).

**Figure 4.**
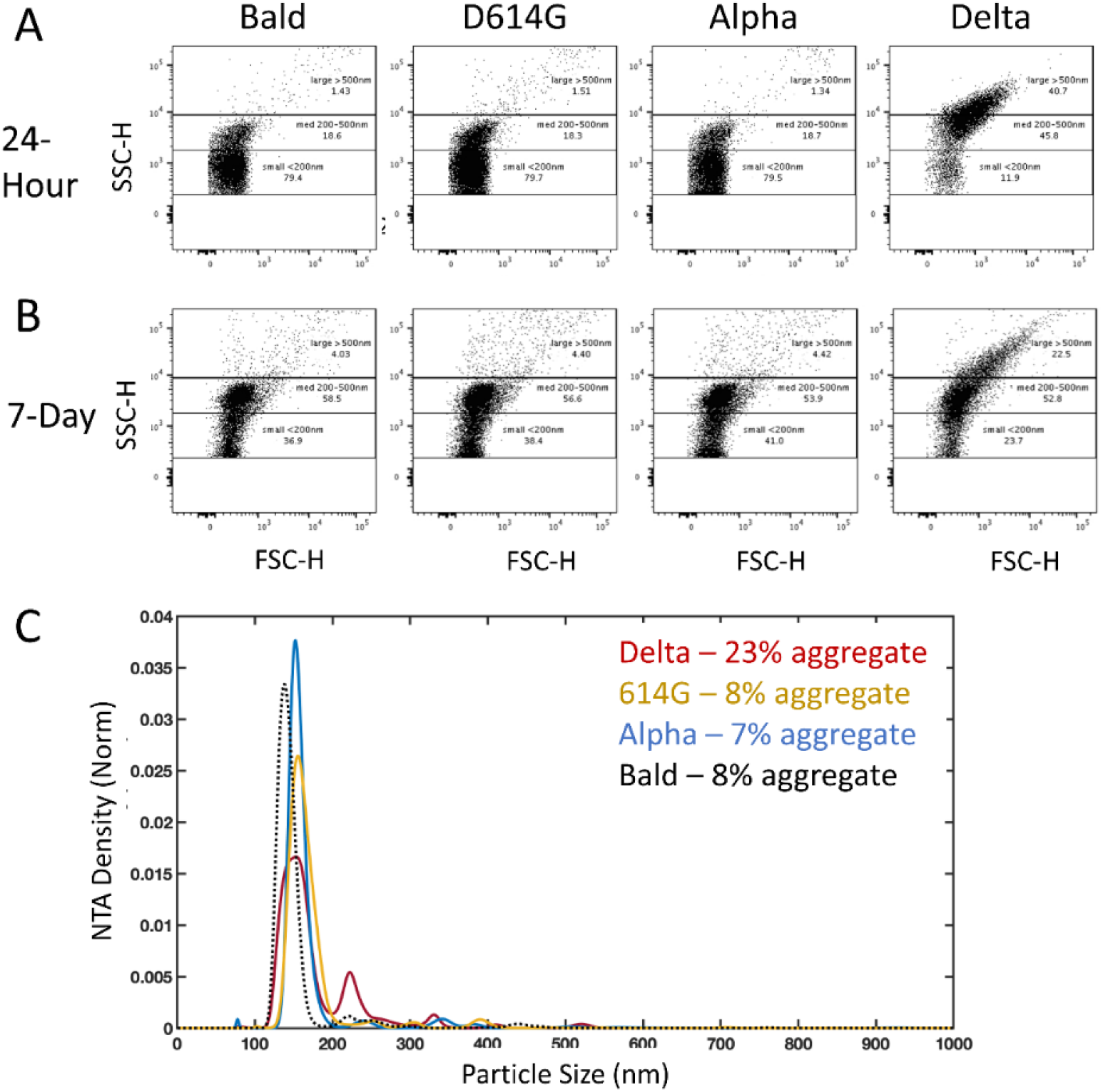
Flow cytometry and NTA detection of Delta PP aggregates. Representative flow cytometry of variant PPs conducted 24-hours (A) and 7-days (B) post-harvest. A population of particles over 200 nm in size are present in the Delta PPs and are minimally (< 10%) represented in the other PP variant samples. (C) NTA of the same samples of PPs at 24-hours post-harvest shows about half as many particles in the single PP range in the Delta sample compared to the other variants, and detection of a larger proportion of PPs in aggregates.

### 3.4 Cryo-electron microscopy of Delta variant PPs

To further evaluate the interactions between spike variant PPs, we used cryo-EM which preserves PPs in hydrated vitreous ice, free from heavy metal stains and potential artifacts of drying that are inherent to negative stain TEM. Delta and all other variant PPs (4 hours and 24 hours post-harvest) were adhered to holey grids and plunge frozen. Imaging confirmed the observations from negative staining that PPs harboring D614G, Alpha, or Omicron BA.1 spikes, as well as the Bald PPs without any spikes, existed mostly as single PPs or in some rare cases in groups of 2-3 closely associated PPs. In contrast, Delta and Delta AY.4.2 PPs mostly existed in aggregates (Figure 5A, B). LDL particles near and far from PPs were observed in all samples with a size range between 19-22 nm. Bald vesicles, damaged PPs and free gag were also seen on occasion. The log-normal cumulative distributions show the shift expected for time-dependent aggregation (Figure 5C). There is no evidence to support a significant difference in aggregate sizes between Delta and Delta AY.4.2.

**Figure 5.**
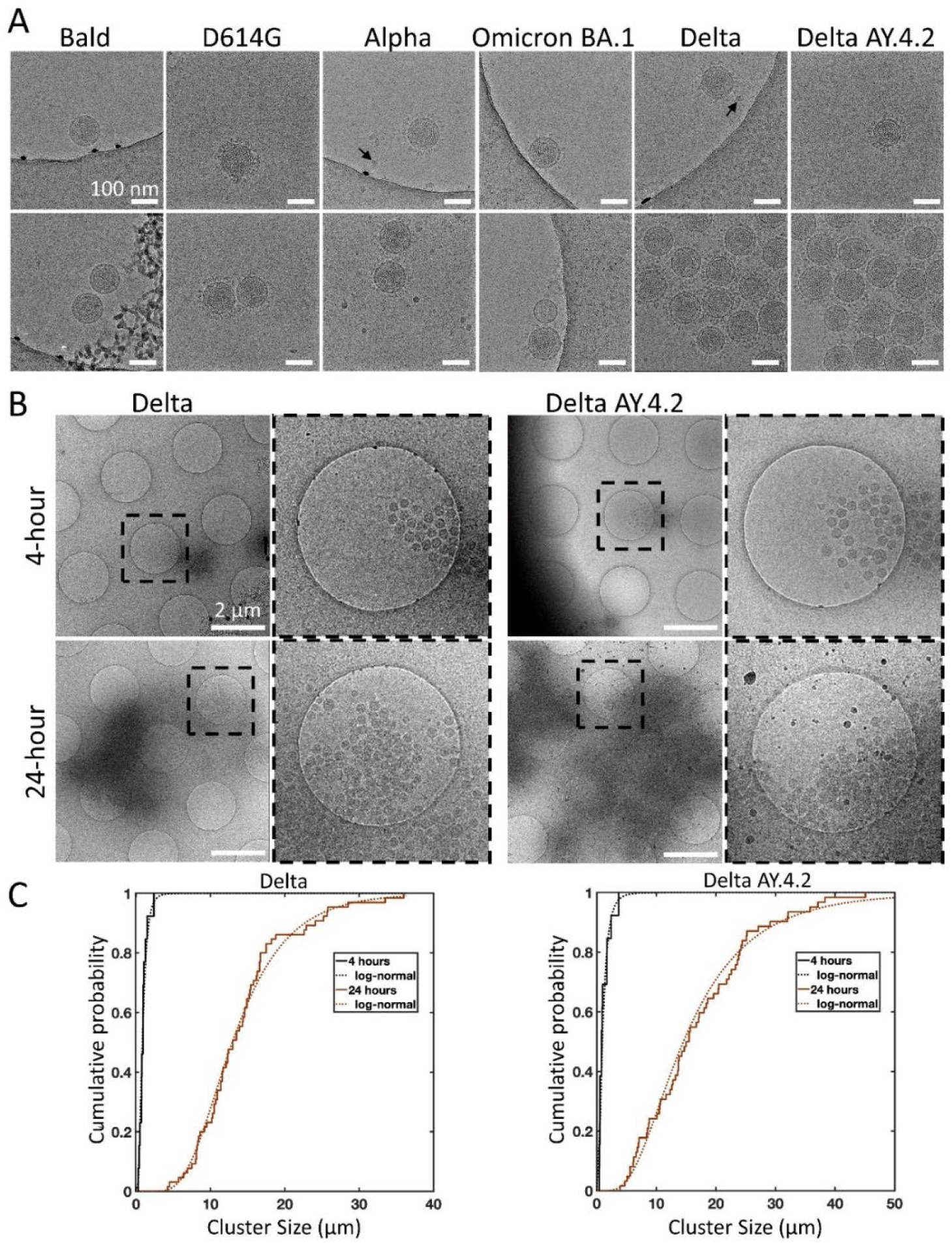
Cryo-electron microscopy of spike variant PPs. (A) Examples of singles (top panel), doubles, or aggregates (lower panel) of each variant PP. Spike variant is identified above. Black arrows indicate lipoproteins. The average PP envelope diameters for each variant are 118.6 (1.1), 118.5 (1.2), 119.6 (0.9), 120.1 (1.3), 115.8 (0.8), 118.4 (0.6) nm for Bald, D614G, Alpha, Omicron BA.1, Delta, and Delta AY.4.2, respectively (mean (SEM), n = 9, 33, 40, 18, 65, and 98. (B) Low magnification overviews of Delta and Delta AY.4.2 PP aggregates prepared 4 or 24 hours post-harvest. Dashed box shows the zoomed-in area. (C) The log-normal cumulative distributions for Delta and Delta AY.4.2 clearly show aggregation over time for both variants. At 4 and 24 hours, Delta aggregate sizes (longest axis) range from 0.28 – 2.39 and 4.20 – 36.0 μm, with log-normal parameters mu (sigma) −0.18 (0.56) and 2.56 (0.44) corresponding to means of 0.97 and 14.25 μm, respectively. There is no evidence for significant differences in aggregate sizes between Delta and Delta AY.4.2 for both 4 and 24 hours.

## 4. DISCUSSION

We demonstrate by four independent techniques—negative stain TEM, flow cytometry, NTA, and cryo-EM— clustering into aggregates of PPs bearing the Delta variant spike or the closely related Delta sublineage Delta AY.4.2 spike, while PPs bearing other spike variants and Bald PPs do not aggregate. Since the PPs were prepared in parallel, handled under identical conditions, and factors that could promote aggregation such as pH changes, freeze-thaw cycles, and high-speed ultracentrifugation were avoided, the observed aggregation of Delta PPs most likely reflects a unique property of the Delta and Delta AY.4.2 spikes. The observation that Delta and Delta AY.4.2 PPs continued to aggregate in solution while stored at 4°C suggests that aggregation occurs after budding from the producer cell, however interaction between PPs could also initiate during biosynthesis and budding from the producer cell.

Unlike SARS-CoV-2 which buds into the ERGIC compartment during final assembly, retroviruses, including MLV, generally bud directly from the plasma membrane (but not always, see [51]). Thus, it is not known when or if Delta variant SARS-CoV-2 aggregates and this is currently being investigated. Delta SARS-CoV-2 could aggregate while budding into the ERGIC, during egress through alkalized lysosomal organelles [52], after egress in the extracellular milieu, or even on the target cell plasma membrane. The ability to aggregate may depend on the concentration of viral particles in each environment. The fact that Delta PP continue to aggregate (Figure 2, 4, and 5) is consistent with a mass action mechanism for Delta PP aggregation. Furthermore, to produce PPs, a 19 aa C-terminal truncated version of each variant spike was expressed, which has been shown to increase the amount of spike incorporated into the PP envelope and PP infectivity [47,48]. Truncations in the cytoplasmic tail could modify properties of the spike ectodomain structure and function [53,54]. Though each variant spike possessed the same truncation, it is not impossible that the truncation uniquely effected the Delta spike ectodomain, conferring the aggregation property.

Examples of Delta PPs with apparent spike tip interactions were observed by negative stain but these interactions did not extend to lateral aggregation of spike proteins on the surface of a PP Ongoing cryo-electron tomography studies will reveal the nature of the interactions between aggregated PPs.

Spike-mediated aggregation differs from antibody-driven aggregation of viral particles expected from polyvalent neutralizing antibodies, as those binding constants are expected to be much stronger. Thus, viral particles would be tightly packed and not likely to disaggregate at the cell surface. The spike tip interactions are more likely to come apart upon surface binding and receptor competition for the RBD at the tip of the spike.

Analysis of the mutations present in the Delta and Delta AY.4.2 spikes compared to other variants may provide clues as to their unique property to aggregate. Delta and Delta AY.4.2 amino acid sequences are similar, except for 4 additional substitutions in Delta AY.4.2 (T95I, Y145H, A222V, and K458R). T95I is also present on the Omicron BA.1 spike and residue Y145 is also substituted in Omicron BA.1 (Y145D). A222V and K458R are unique to Delta AY.4.2. As the presence of these mutations in Delta AY.4.2 do not abrogate or enhance the aggregation of Delta AY.4.2 compared to Delta, they appear to have no effect on aggregation.

Most of the other mutations on the Delta and Delta AY.4.2 spikes are shared with other variants. The D614G mutation is present in all variants, substitution L452R located in the RBD is present in the Kappa variant (B.1.617.1), and a similar substitution L452Q occurs in the Lambda variant (C.37). A second RBD substitution at T478K, is also found in Omicron BA.1 and BA.2. The mutation P681R is present in Kappa, and P681H exists in Alpha, Omicron BA.1, BA.2 and Mu (B.1.621). Finally, substitution mutation D950N is also shared with Mu. Because the non-Delta variants studied here do not aggregate, it is unlikely that any of their Delta-shared mutations can be aggregation-dominant.

There are, however, three residues, E156, F157 and R158, in the NTD of Delta and Delta AY.4.2 that are uniquely and identically mutated: substitution E156G, and deletions at F157 and R158. It is possible that these three mutations in the NTD are sufficient to bestow the aggregation property, or perhaps the unique combination of mutations occurring on the Delta spike drive aggregation synergistically.

The clustering of Delta PPs could account for the faster and larger initial infection observed in entry assays with the Delta PPs [35]. Because the number of spike trimers is larger on an aggregate comprising multiple PPs, and the cell surface contact area is larger for any collision between the aggregate and a target cell, the effective on-rate for aggregate binding should be larger, resulting in faster binding. Furthermore, the avidity of the aggregate to the target cell would be enhanced manyfold due to the multiple potential binding partners on a single contacting surface. Moreover, the increased dwell time at that contact area will allow for diffusional and conformational motions of proteins and lipids to increase the chance of membrane fusion, as these factors are important for avoiding hemifusion and promoting full fusion [55]. All these factors should lead to the relatively higher initial rate of PP entry into target cells from aggregated PPs. Whether or not aggregates could enable the simultaneous delivery of multiple copies of entry reporter genes to target cell is not clear, since the PPs need not display receptor and may not fuse to each other, even in an endosome, so one entry event may or may not mean copies of genes enter. Implicitly, there would be more overall binding events for unaggregated PPs, each at another site. However, if the probability for PP entry was low due to unbinding, then the factors discussed above to increase PP avidity would tend to increase overall fusion and its rate.

In summary, an ultrastructural analysis of retrovirus pseudotyped particles with coronavirus spike proteins from variants of SARS-CoV-2 led to a serendipitous discovery of significant aggregation when the Delta variant of the spike is expressed, but not upon expression of three other variants. Viral aggregation can impart fitness benefits by protecting virions from environmental hazards and by effecting simultaneous delivery of multiple viral genomes, or collective infection [56]. Notably, collective infection can favor initial infection in some contexts [57]. Likely, the size and number of virions per aggregate is important to increasing infectivity. Too large and it would effectively reduce infectious units below a threshold. Too few virions in an aggregate and the benefit of collective infection is not gained. The unique property of the Delta spike to aggregate PPs may underlie the faster infection by Delta PPs. Furthermore, spike mediated aggregation could be part of the molecular mechanism by which Delta variant SARS-CoV-2 achieves increased transmissibility and faster infection with a higher viral load. The continued clustering over time indicates that the underlying factor for Delta PP clustering may be the spike tip interactions of the spike protein, which in turn may indicate an adhesivity of the viral surface recognized by the immune system thus altering the balance of host antiviral response towards inflammation.

## Author Contributions

Conceptualization, J.D.P., J.Z.; methodology, J.D.P., W.F., D.M., F.Z., P.B.S; formal analysis, J.D.P., P.S.B., W.F., D.M., F.Z.; investigation, J.D.P., W.F., D.M., F.Z.; writing—original draft preparation, J.D.P., J.Z.; writing—review and editing, J.D.P., J.Z., D.M., F.Z., P.S.B., W.F., J.L.; visualization, J.D.P., D.M., F.Z., P.S.B.; supervision, J.Z.

All authors have read and agreed to the published version of the manuscript

## Institutional Review Board Statement

Not applicable

## Informed Consent Statement

Not applicable

## Acknowledgments

We thank Rick Kuo-Jui Huang from the National Cancer Institute and Stéphane Mahé from Thermo Fisher Scientific for their technical support at the FEI Tecnai T20 electron microscope.

## References

1. Plante, J.A.; Mitchell, B.M.; Plante, K.S.; Debbink, K.; Weaver, S.C.; Menachery, V.D. The variant gambit: COVID-19’s next move. Cell Host Microbe 2021, 29, 508–515, doi:10.1016/j.chom.2021.02.020.

2. Fehr, A.R.; Perlman, S. Coronaviruses: An Overview of Their Replication and Pathogenesis. Methods Mol Biol 2015, 1282, 1–23, doi:10.1007/978-1-4939-2438-7_1.

3. Bosch, B.J.; van der Zee, R.; de Haan, C.A.; Rottier, P.J. The coronavirus spike protein is a class I virus fusion protein: structural and functional characterization of the fusion core complex. J Virol 2003, 77, 8801–8811, doi:10.1128/jvi.77.16.8801-8811.2003.

4. Li, F. Structure, Function, and Evolution of Coronavirus Spike Proteins. Annu Rev Virol 2016, 3, 237–261, doi:10.1146/annurev-virology-110615-042301.

5. Walls, A.C.; Park, Y.J.; Tortorici, M.A.; Wall, A.; McGuire, A.T.; Veesler, D. Structure, Function, and Antigenicity of the SARS-CoV-2 Spike Glycoprotein. Cell 2020, 183, 1735, doi:10.1016/j.cell.2020.11.032.

6. Wrapp, D.; Wang, N.; Corbett, K.S.; Goldsmith, J.A.; Hsieh, C.L.; Abiona, O.; Graham, B.S.; McLellan, J.S. Cryo-EM structure of the 2019-nCoV spike in the prefusion conformation. Science 2020, 367, 1260–1263, doi:10.1126/science.abb2507.

7. Watanabe, Y.; Allen, J.D.; Wrapp, D.; McLellan, J.S.; Crispin, M. Site-specific glycan analysis of the SARS-CoV-2 spike. Science 2020, 369, 330–333, doi:10.1126/science.abb9983.

8. Kumar, S.; Maurya, V.K.; Prasad, A.K.; Bhatt, M.L.B.; Saxena, S.K. Structural, glycosylation and antigenic variation between 2019 novel coronavirus (2019-nCoV) and SARS coronavirus (SARS-CoV). Virusdisease 2020, 31, 13–21, doi:10.1007/s13337-020-00571-5.

9. Hoffmann, M.; Kleine-Weber, H.; Schroeder, S.; Kruger, N.; Herrler, T.; Erichsen, S.; Schiergens, T.S.; Herrler, G.; Wu, N.H.; Nitsche, A.; et al. SARS-CoV-2 Cell Entry Depends on ACE2 and TMPRSS2 and Is Blocked by a Clinically Proven Protease Inhibitor. Cell 2020, 181, 271–280 e278, doi:10.1016/j.cell.2020.02.052.

10. Ke, Z.; Oton, J.; Qu, K.; Cortese, M.; Zila, V.; McKeane, L.; Nakane, T.; Zivanov, J.; Neufeldt, C.J.; Cerikan, B.; et al. Structures and distributions of SARS-CoV-2 spike proteins on intact virions. Nature 2020, 588, 498–502, doi:10.1038/s41586-020-2665-2.

11. Turonova, B.; Sikora, M.; Schurmann, C.; Hagen, W.J.H.; Welsch, S.; Blanc, F.E.C.; von Bulow, S.; Gecht, M.; Bagola, K.; Horner, C.; et al. In situ structural analysis of SARS-CoV-2 spike reveals flexibility mediated by three hinges. Science 2020, 370, 203–208, doi:10.1126/science.abd5223.

12. Choi, Y.K.; Cao, Y.; Frank, M.; Woo, H.; Park, S.J.; Yeom, M.S.; Croll, T.I.; Seok, C.; Im, W. Structure, Dynamics, Receptor Binding, and Antibody Binding of the Fully Glycosylated Full-Length SARS-CoV-2 Spike Protein in a Viral Membrane. J Chem Theory Comput 2021, 17, 2479–2487, doi:10.1021/acs.jctc.0c01144.

13. Peacock, T.P.; Goldhill, D.H.; Zhou, J.; Baillon, L.; Frise, R.; Swann, O.C.; Kugathasan, R.; Penn, R.; Brown, J.C.; Sanchez-David, R.Y.; et al. The furin cleavage site in the SARS-CoV-2 spike protein is required for transmission in ferrets. Nat Microbiol 2021, 6, 899–909, doi:10.1038/s41564-021-00908-w.

14. Tang, T.; Bidon, M.; Jaimes, J.A.; Whittaker, G.R.; Daniel, S. Coronavirus membrane fusion mechanism offers a potential target for antiviral development. Antiviral Res 2020, 178, 104792, doi:10.1016/j.antiviral.2020.104792.

15. Walls, A.C.; Tortorici, M.A.; Snijder, J.; Xiong, X.; Bosch, B.J.; Rey, F.A.; Veesler, D. Tectonic conformational changes of a coronavirus spike glycoprotein promote membrane fusion. Proc Natl Acad Sci U S A 2017, 114, 11157–11162, doi:10.1073/pnas.1708727114.

16. Millet, J.K.; Whittaker, G.R. Host cell entry of Middle East respiratory syndrome coronavirus after two-step, furin-mediated activation of the spike protein. Proc Natl Acad Sci U S A 2014, 111, 15214–15219, doi:10.1073/pnas.1407087111.

17. Zhou, P.; Yang, X.L.; Wang, X.G.; Hu, B.; Zhang, L.; Zhang, W.; Si, H.R.; Zhu, Y.; Li, B.; Huang, C.L.; et al. A pneumonia outbreak associated with a new coronavirus of probable bat origin. Nature 2020, 579, 270–273, doi:10.1038/s41586-020-2012-7.

18. Zhu, N.; Zhang, D.; Wang, W.; Li, X.; Yang, B.; Song, J.; Zhao, X.; Huang, B.; Shi, W.; Lu, R.; et al. A Novel Coronavirus from Patients with Pneumonia in China, 2019. N Engl J Med 2020, 382, 727–733, doi:10.1056/NEJMoa2001017.

19. Gobeil, S.M.; Janowska, K.; McDowell, S.; Mansouri, K.; Parks, R.; Stalls, V.; Kopp, M.F.; Manne, K.; Li, D.; Wiehe, K.; et al. Effect of natural mutations of SARS-CoV-2 on spike structure, conformation, and antigenicity. Science 2021, 373, doi:10.1126/science.abi6226.

20. Negi, S.S.; Schein, C.H.; Braun, W. Regional and temporal coordinated mutation patterns in SARS-CoV-2 spike protein revealed by a clustering and network analysis. Sci Rep 2022, 12, 1128, doi:10.1038/s41598-022-04950-4.

21. Konings, F.; Perkins, M.D.; Kuhn, J.H.; Pallen, M.J.; Alm, E.J.; Archer, B.N.; Barakat, A.; Bedford, T.; Bhiman, J.N.; Caly, L.; et al. SARS-CoV-2 Variants of Interest and Concern naming scheme conducive for global discourse. Nat Microbiol 2021, 6, 821–823, doi:10.1038/s41564-021-00932-w.

22. Li, Q.; Wu, J.; Nie, J.; Zhang, L.; Hao, H.; Liu, S.; Zhao, C.; Zhang, Q.; Liu, H.; Nie, L.; et al. The Impact of Mutations in SARS-CoV-2 Spike on Viral Infectivity and Antigenicity. Cell 2020, 182, 1284–1294 e1289, doi:10.1016/j.cell.2020.07.012.

23. Zhang, J.; Cai, Y.; Xiao, T.; Lu, J.; Peng, H.; Sterling, S.M.; Walsh, R.M.; Rits-Volloch, S.; Sliz, P.; Chen, B. Structural impact on SARS-CoV-2 spike protein by D614G substitution. bioRxiv 2020, doi:10.1101/2020.10.13.337980.

24. Yurkovetskiy, L.; Wang, X.; Pascal, K.E.; Tomkins-Tinch, C.; Nyalile, T.P.; Wang, Y.; Baum, A.; Diehl, W.E.; Dauphin, A.; Carbone, C.; et al. Structural and Functional Analysis of the D614G SARS-CoV-2 Spike Protein Variant. Cell 2020, 183, 739–751 e738, doi:10.1016/j.cell.2020.09.032.

25. Zhang, L.; Jackson, C.B.; Mou, H.; Ojha, A.; Peng, H.; Quinlan, B.D.; Rangarajan, E.S.; Pan, A.; Vanderheiden, A.; Suthar, M.S.; et al. SARS-CoV-2 spike-protein D614G mutation increases virion spike density and infectivity. Nat Commun 2020, 11, 6013, doi:10.1038/s41467-020-19808-4.

26. Rambaut, A.; Holmes, E.C.; O’Toole, A.; Hill, V.; McCrone, J.T.; Ruis, C.; du Plessis, L.; Pybus, O.G. A dynamic nomenclature proposal for SARS-CoV-2 lineages to assist genomic epidemiology. Nat Microbiol 2020, 5, 1403–1407, doi:10.1038/s41564-020-0770-5.

27. Wang, C.; Liu, Z.; Chen, Z.; Huang, X.; Xu, M.; He, T.; Zhang, Z. The establishment of reference sequence for SARS-CoV-2 and variation analysis. J Med Virol 2020, 92, 667–674, doi:10.1002/jmv.25762.

28. Korber, B.; Fischer, W.M.; Gnanakaran, S.; Yoon, H.; Theiler, J.; Abfalterer, W.; Hengartner, N.; Giorgi, E.E.; Bhattacharya, T.; Foley, B.; et al. Tracking Changes in SARS-CoV-2 Spike: Evidence that D614G Increases Infectivity of the COVID-19 Virus. Cell 2020, 182, 812–827 e819, doi:10.1016/j.cell.2020.06.043.

29. Lubinski, B.; Fernandes, M.H.V.; Frazier, L.; Tang, T.; Daniel, S.; Diel, D.G.; Jaimes, J.A.; Whittaker, G.R. Functional evaluation of the P681H mutation on the proteolytic activation of the SARS-CoV-2 variant B.1.1.7 (Alpha) spike. iScience 2022, 25, 103589, doi:10.1016/j.isci.2021.103589.

30. Meng, B.; Kemp, S.A.; Papa, G.; Datir, R.; Ferreira, I.; Marelli, S.; Harvey, W.T.; Lytras, S.; Mohamed, A.; Gallo, G.; et al. Recurrent emergence of SARS-CoV-2 spike deletion H69/V70 and its role in the Alpha variant B.1.1.7. Cell Rep 2021, 35, 109292, doi:10.1016/j.celrep.2021.109292.

31. Starr, T.N.; Greaney, A.J.; Hilton, S.K.; Ellis, D.; Crawford, K.H.D.; Dingens, A.S.; Navarro, M.J.; Bowen, J.E.; Tortorici, M.A.; Walls, A.C.; et al. Deep Mutational Scanning of SARS-CoV-2 Receptor Binding Domain Reveals Constraints on Folding and ACE2 Binding. Cell 2020, 182, 1295–1310 e1220, doi:10.1016/j.cell.2020.08.012.

32. Grabowski, F.; Preibisch, G.; Gizinski, S.; Kochanczyk, M.; Lipniacki, T. SARS-CoV-2 Variant of Concern 202012/01 Has about Twofold Replicative Advantage and Acquires Concerning Mutations. Viruses 2021, 13, doi:10.3390/v13030392.

33. Singh, J.; Rahman, S.A.; Ehtesham, N.Z.; Hira, S.; Hasnain, S.E. SARS-CoV-2 variants of concern are emerging in India. Nat Med 2021, 27, 1131–1133, doi:10.1038/s41591-021-01397-4.

34. Dhar, M.S.; Marwal, R.; Vs, R.; Ponnusamy, K.; Jolly, B.; Bhoyar, R.C.; Sardana, V.; Naushin, S.; Rophina, M.; Mellan, T.A.; et al. Genomic characterization and epidemiology of an emerging SARS-CoV-2 variant in Delhi, India. Science 2021, 374, 995–999, doi:10.1126/science.abj9932.

35. Zhang, J.; Xiao, T.; Cai, Y.; Lavine, C.L.; Peng, H.; Zhu, H.; Anand, K.; Tong, P.; Gautam, A.; Mayer, M.L.; et al. Membrane fusion and immune evasion by the spike protein of SARS-CoV-2 Delta variant. Science 2021, eabl9463, doi:10.1126/science.abl9463.

36. Teyssou, E.; Delagreverie, H.; Visseaux, B.; Lambert-Niclot, S.; Brichler, S.; Ferre, V.; Marot, S.; Jary, A.; Todesco, E.; Schnuriger, A.; et al. The Delta SARS-CoV-2 variant has a higher viral load than the Beta and the historical variants in nasopharyngeal samples from newly diagnosed COVID-19 patients. J Infect 2021, 83, e1–e3, doi:10.1016/j.jinf.2021.08.027.

37. Bolze, A.; Cirulli, E.T.; Luo, S.; White, S.; Wyman, D.; Rossi, A.D.; Machado, H.; Cassens, T.; Jacobs, S.; Schiabor Barrett, K.M.; et al. SARS-CoV-2 variant Delta rapidly displaced variant Alpha in the United States and led to higher viral loads. medRxiv 2021, 2021.2006.2020.21259195, doi:10.1101/2021.06.20.21259195.

38. Twohig, K.A.; Nyberg, T.; Zaidi, A.; Thelwall, S.; Sinnathamby, M.A.; Aliabadi, S.; Seaman, S.R.; Harris, R.J.; Hope, R.; Lopez-Bernal, J.; et al. Hospital admission and emergency care attendance risk for SARS-CoV-2 delta (B.1.617.2) compared with alpha (B.1.1.7) variants of concern: a cohort study. Lancet Infect Dis 2022, 22, 35–42, doi:10.1016/S1473-3099(21)00475-8.

39. McCallum, M.; Bassi, J.; De Marco, A.; Chen, A.; Walls, A.C.; Di Iulio, J.; Tortorici, M.A.; Navarro, M.J.; Silacci-Fregni, C.; Saliba, C.; et al. SARS-CoV-2 immune evasion by the B.1.427/B.1.429 variant of concern. Science 2021, 373, 648–654, doi:10.1126/science.abi7994.

40. Neerukonda, S.N.; Vassell, R.; Lusvarghi, S.; Wang, R.; Echegaray, F.; Bentley, L.; Eakin, A.E.; Erlandson, K.J.; Katzelnick, L.C.; Weiss, C.D.; et al. SARS-CoV-2 Delta Variant Displays Moderate Resistance to Neutralizing Antibodies and Spike Protein Properties of Higher Soluble ACE2 Sensitivity, Enhanced Cleavage and Fusogenic Activity. Viruses 2021, 13, doi:10.3390/v13122485.

41. Planas, D.; Veyer, D.; Baidaliuk, A.; Staropoli, I.; Guivel-Benhassine, F.; Rajah, M.M.; Planchais, C.; Porrot, F.; Robillard, N.; Puech, J.; et al. Reduced sensitivity of SARS-CoV-2 variant Delta to antibody neutralization. Nature 2021, 596, 276–280, doi:10.1038/s41586-021-03777-9.

42. Saito, A.; Irie, T.; Suzuki, R.; Maemura, T.; Nasser, H.; Uriu, K.; Kosugi, Y.; Shirakawa, K.; Sadamasu, K.; Kimura, I.; et al. Enhanced fusogenicity and pathogenicity of SARS-CoV-2 Delta P681R mutation. Nature 2022, 602, 300–306, doi:10.1038/s41586-021-04266-9.

43. Mlcochova, P.; Kemp, S.A.; Dhar, M.S.; Papa, G.; Meng, B.; Ferreira, I.; Datir, R.; Collier, D.A.; Albecka, A.; Singh, S.; et al. SARS-CoV-2 B.1.617.2 Delta variant replication and immune evasion. Nature 2021, 599, 114–119, doi:10.1038/s41586-021-03944-y.

44. Motozono, C.; Toyoda, M.; Zahradnik, J.; Saito, A.; Nasser, H.; Tan, T.S.; Ngare, I.; Kimura, I.; Uriu, K.; Kosugi, Y.; et al. SARS-CoV-2 spike L452R variant evades cellular immunity and increases infectivity. Cell Host Microbe 2021, 29, 1124–1136 e1111, doi:10.1016/j.chom.2021.06.006.

45. Millet, J.K.; Whittaker, G.R. Murine Leukemia Virus (MLV)-based Coronavirus Spike-pseudotyped Particle Production and Infection. Bio Protoc 2016, 6, doi:10.21769/BioProtoc.2035.

46. Xu, M.; Pradhan, M.; Gorshkov, K.; Petersen, J.D.; Shen, M.; Guo, H.; Zhu, W.; Klumpp-Thomas, C.; Michael, S.; Itkin, M.; et al. A high throughput screening assay for inhibitors of SARS-CoV-2 pseudotyped particle entry. SLAS Discov 2022, 27, 86–94, doi:10.1016/j.slasd.2021.12.005.

47. Johnson, M.C.; Lyddon, T.D.; Suarez, R.; Salcedo, B.; LePique, M.; Graham, M.; Ricana, C.; Robinson, C.; Ritter, D.G. Optimized Pseudotyping Conditions for the SARS-COV-2 Spike Glycoprotein. J Virol 2020, 94, doi:10.1128/JVI.01062-20.

48. Giroglou, T.; Cinatl, J., Jr.; Rabenau, H.; Drosten, C.; Schwalbe, H.; Doerr, H.W.; von Laer, D. Retroviral vectors pseudotyped with severe acute respiratory syndrome coronavirus S protein. J Virol 2004, 78, 9007–9015, doi:10.1128/JVI.78.17.9007-9015.2004.

49. Mastronarde, D.N. Automated electron microscope tomography using robust prediction of specimen movements. J Struct Biol 2005, 152, 36–51, doi:10.1016/j.jsb.2005.07.007.

50. Yeager, M.; Wilson-Kubalek, E.M.; Weiner, S.G.; Brown, P.O.; Rein, A. Supramolecular organization of immature and mature murine leukemia virus revealed by electron cryo-microscopy: implications for retroviral assembly mechanisms. Proc Natl Acad Sci U S A 1998, 95, 7299–7304, doi:10.1073/pnas.95.13.7299.

51. Houzet, L.; Gay, B.; Morichaud, Z.; Briant, L.; Mougel, M. Intracellular assembly and budding of the Murine Leukemia Virus in infected cells. Retrovirology 2006, 3, 12, doi:10.1186/1742-4690-3-12.

52. Ghosh, S.; Dellibovi-Ragheb, T.A.; Kerviel, A.; Pak, E.; Qiu, Q.; Fisher, M.; Takvorian, P.M.; Bleck, C.; Hsu, V.W.; Fehr, A.R.; et al. beta-Coronaviruses Use Lysosomes for Egress Instead of the Biosynthetic Secretory Pathway. Cell 2020, 183, 1520–1535 e1514, doi:10.1016/j.cell.2020.10.039.

53. Cathomen, T.; Naim, H.Y.; Cattaneo, R. Measles viruses with altered envelope protein cytoplasmic tails gain cell fusion competence. J Virol 1998, 72, 1224–1234, doi:10.1128/JVI.72.2.1224-1234.1998.

54. Zingler, K.; Littman, D.R. Truncation of the cytoplasmic domain of the simian immunodeficiency virus envelope glycoprotein increases env incorporation into particles and fusogenicity and infectivity. J Virol 1993, 67, 2824–2831, doi:10.1128/JVI.67.5.2824-2831.1993.

55. Chernomordik, L.V.; Frolov, V.A.; Leikina, E.; Bronk, P.; Zimmerberg, J. The pathway of membrane fusion catalyzed by influenza hemagglutinin: restriction of lipids, hemifusion, and lipidic fusion pore formation. J Cell Biol 1998, 140, 1369–1382, doi:10.1083/jcb.140.6.1369.

56. Sanjuan, R.; Thoulouze, M.I. Why viruses sometimes disperse in groups?(dagger). Virus Evol 2019, 5, vez014, doi:10.1093/ve/vez014.

57. Andreu-Moreno, I.; Sanjuan, R. Collective Infection of Cells by Viral Aggregates Promotes Early Viral Proliferation and Reveals a Cellular-Level Allee Effect. Curr Biol 2018, 28, 3212–3219 e3214, doi:10.1016/j.cub.2018.08.028.

